# Ultradian regulation of rest and activity bouts in mice

**DOI:** 10.1101/836478

**Authors:** Bharath Ananthasubramaniam, Johanna H. Meijer

## Abstract

The suprachiasmatic nucleus (SCN), which serves as the central pacemaker in mammals, regulates the 24-hour rhythm in behavioral activity. However, it is currently unclear whether and how bouts of activity and rest are regulated within the 24-hour cycle (i.e., over ultradian time scales). Therefore, we used passive infrared sensors to measure behavior in mice housed under either a light-dark (LD) cycle or continuous darkness (DD). We found that a probabilistic Markov model captures the ultradian changes in the behavioral state over a 24-hour cycle. In this model, the animal’s behavioral state in the next time interval is determined solely by the animal’s current behavioral state and by the “toss” of a proverbial “biased coin”. We found that the bias of this “coin” is regulated by light input and by the phase of the clock. Moreover, the bias of this “coin” for an animal is related to the average length of rest and activity bouts in that animal. In LD conditions, the average length of rest bouts was greater during the day compared to during the night, whereas the average length of activity bouts was greater during the night compared to during the day. Importantly, we also found that day-night changes in the rest bout lengths were significantly greater than day-night changes in the activity bout lengths. Finally, in DD conditions, the activity and rest bouts also differed between subjective night and subjective day, albeit to a lesser extent compared to LD conditions. The persistent differences in bout length over the circadian cycle following loss of the external LD cycle indicate that the central pacemaker plays a role in regulating rest and activity bouts on an ultradian time scale.

## 1 Introduction

In most organisms, the circadian clock facilitates adaptation to the natural periodic light cycle. This clock regulates a wide range of physiological processes, including behavior (Herzog, 2007). Therefore, behavior has been used to determine the state of the clock *in vivo* since the early days of the field of chronobiology (Pittendrigh, 1960; Pittendrigh and Daan, 1976). In mammals, the circadian clock is located in the suprachiasmatic nucleus (SCN) at the base of the hypothalamus (Ralph et al., 1990). The neurons in the SCN have near 24-hour oscillations in both protein expression and neuronal firing (Nakamura et al., 2002; Quintero et al., 2003; Schaap et al., 2003; Yamaguchi et al., 2003; Hastings et al., 2018).

Recording the frequency of action potential firing in the SCN of freely moving animals has allowed researchers to measure the degree of correspondence between SCN firing and behavioral activity (Houben et al., 2009, 2014). These studies showed that the onset and offset of behavioral activity are regulated probabilistically by differences in firing between high levels of firing activity during the day and low levels during the night (Houben et al., 2009). Moreover, the waveform of the SCN’s firing patterns is correlated with the distribution of behavioral activity within the active phase (Houben et al., 2009, 2014). However, whether—and to what extent—temporal behavior is regulated within the circadian cycle (i.e., over ultradian time scales) is largely unknown.

Here, we examined whether bouts of activity and rest (i.e., prolonged stretches of activity and rest, respectively) are regulated at ultradian time scales in mice. We fit a simple probabilistic model of the transitions between behavioral states to behavioral data collected under a light-dark (LD) cycle or continuous darkness (DD). Our model shows how bouts of rest and activity are regulated on a scale of seconds to minutes. In addition, the model shows that changes in the duration of rest bouts, rather than changes in the duration of activity bouts, determine the differences in activity between day and night.

## 2 Materials and Methods

### 2.1 Ethics statement

All animal experiments were performed in accordance with Dutch law and were approved by the Animal Ethics Committee at Leiden University Medical Center (Leiden, the Netherlands).

### 2.2 Animals

Wild-type C57BL6 mice were purchased from Harlan (Harlan, Horst, the Netherlands). All mice were between 12-24 weeks of age.

### 2.3 Behavioral data

Each animal’s behavioral activity was recorded using passive IR (PIR) motion detection sensors (Hygrosens Instruments) mounted under the lid of the cage and connected to a ClockLab data collection system (Actimetrics Software), which recorded sensor activity in 10-sec bins.

Mice were housed under either continuous darkness (DD) or an LD cycle with a 22-hour, 24-hour, or 26-hour period with equal duration of light and dark (also termed T-cycles); for example, a 22-hour T-cycle consisted of 11 hours of light and 11 hours of darkness. Only recordings of mice with at least two circadian cycles in DD or four cycles in an LD cycle were included in our analysis. Furthermore, all mice housed under an LD cycle were entrained to the external *Zeitgeber*. In this study, a total of 17 mice were housed under DD conditions, and 32 mice were housed under a 22-hour (N=8 mice), 24-hour (N=16 mice), or a 26-hour (N=8 mice) T-cycle.

The data consist of the start time of the 10s bin, with lights “on” or “off” marked as “L” and “D”, respectively (Figure 1); in addition, behavioral activity was counted in 10-sec intervals. For this study, activity counts were converted to either “A” (active; activity counts >0) or “R” (rest; activity counts = 0); thus, we studied the duration of activity and rest, not the intensity of activity.

**Figure 1:**
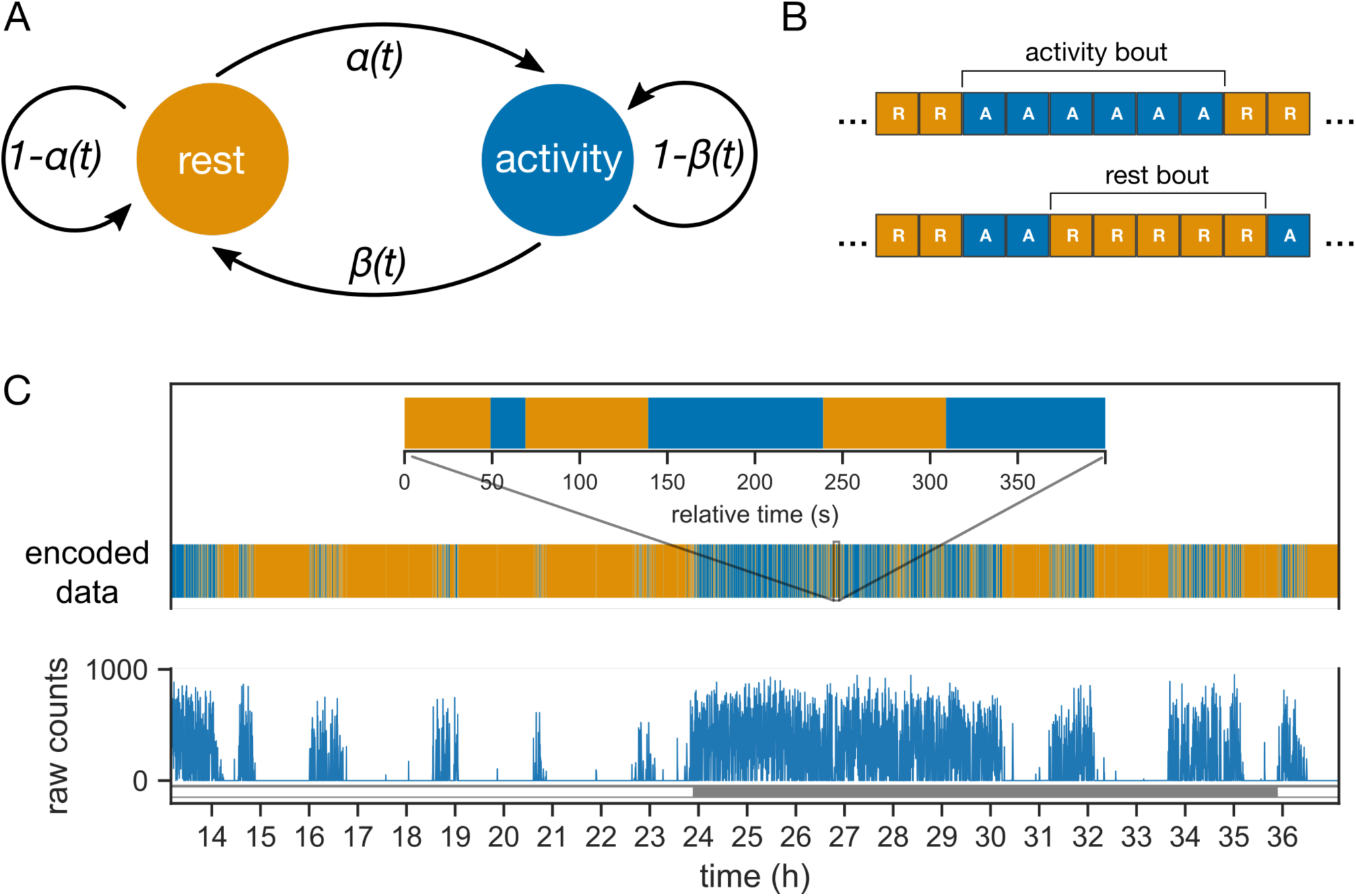
Probabilistic model of mouse behavior. (A) The transitions between the two behavioral states “rest” and “activity” are controlled by the two probability parameters, ***α*** and ***β***. The parameters can vary between day and night (or subjective day and night in constant darkness). (B) The behavioral activity data is encoded into sequences of “activity” (behavioral activity>0) and “rest” (behavioral activity = 0) bins. The encoded sequences are inputs to the model in (A). One or more contiguous occurrences of “activity” and “rest” bins are termed activity and rest bouts, respectively. (C) The behavioral activity of an animal is recorded as raw counts in 10s bins (shown here for a mouse under 24h light-dark cycles). The corresponding encoded sequence is shown with the same color-coding in (B) for rest and activity. Inset zooms into a 400s interval during the night (active phase).

### 2.4 Description of the probabilistic model

The probabilistic model describes the transitions in the animal’s behavioral state between “rest” and “activity” (Figure 1A). The animal’s behavioral state in the next 10-sec bin (*S*_*n*+1_) is determined solely by the behavioral state in the current 10-sec bin (*S*_*n*_) and the probability of transition; such a property defines a Markov model. The transition probability from rest to activity is defined as *α*, and the transition probability from activity to rest is defined as *β*. Probabilities were allowed to change between day and night under LD conditions and between subjective day and subjective night under DD conditions. Under an LD cycle, both the central clock and the external LD cycle contribute to the behavioral state; in contrast, under DD, the effect of the external LD cycle is absent.

For mice in LD, *α* and *β* were fit separately for day and night using the following equations:

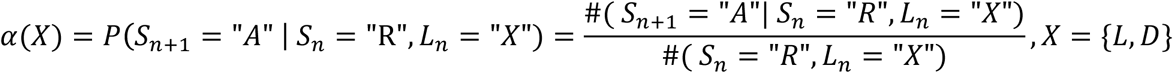

and

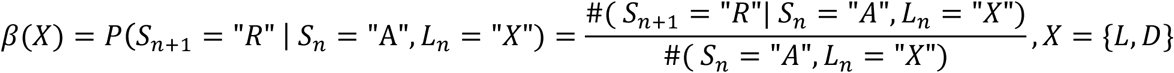

where *L*_*n*_ represents the lighting condition during state *S*_*n*_ and #(·) is the number of occurrences of the condition within the parenthesis. Thus, the data obtained from each animal in LD yields four probabilities, two for the day phase and two for night phase.

For mice housed in DD, cosine curves were fit to the raw behavioral activity counts in order to identify the subjective day and subjective night phases. *α* and *β* were computed for the subjective day and subjective night using the equations shown above, producing a similar set of four probabilities.

We define bouts of activity and rest to be one or more adjacent bins of activity and rest, respectively (Figure 1B). Bouts of activity and rest (Figure 1C) are easier to interpret and identify in the data compared to the probability parameters. The Markov model leads to a geometric distribution of the bout durations, and the mean bout durations are conveniently dependent only upon *α* and *β*. The mean activity bouts and rest bouts (expressed in min) are 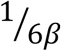 and 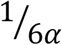. In addition, the average activity during a phase of the clock is defined using the following formula: 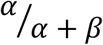.

### 2.5 Statistical analysis

The average activity of the mice was analyzed using the analysis of variance (`aov`) function in R (version 3.5.1). The duration of bouts estimated using the model were first transformed in order to ensure uniform variance across groups; because each animal contributed multiple estimates, they were then analyzed using linear mixed models with the `lmer` function in the R package (“lme4” version 1.1-21).

## 3 Results

### 3.1 Model fits are robust and consistent with the assumptions

First, we determined the ability of our model to produce reasonable parameter estimates that are consistent with the model’s assumptions.

The estimates of transition probability obtained from the behavioral data were stable over the course of acquisition. The estimates of *α* and *β* obtained from the first half of each acquisition were highly correlated with the estimates obtained from the second half (Figure S1A, *α*: r(96)=0.85, p<.001; *β*: r(96)=0.69, p<.001). Thus, the data can be considered stationary for the purposes of this model.

The data also support the Markov assumption made in the probabilistic model, which can be paraphrased as the “the future is independent of the past, given the present”. Estimates derived from the data regarding dependence (via mutual information) between the next state (*S*_*n*+1_) and the previous state (*S*_*n*−1_), given the current state (*S*_*n*_), were all close to zero (Figure S1B), indicating near independence. In summary, our model provides a consistent representation of the data and produces robust estimates of the average duration of activity and rest bouts.

### 3.2 Activity and rest in mice are not restricted to night and day, respectively

Next, we examined the distribution of activity during the day and night under 22-hour, 24-hour, and 26-hour T-cycles (i.e., LD cycles consisting of 11 hours light/11 hours dark, 12 hours light/12 hours dark, and 13 hours light/13 hours dark, respectively).

We defined *average activity* as the average fraction of time an animal was active in an interval; the interval is the length of the day for the average activity during the day. The average activity across day and night (the interval here is the T-cycle period) was similar between the 22-hour and the 24-hour T-cycles, but was significantly higher in the 26-hour T-cycle compared to the 24-hour T-cycle (Figure S2, *F*(2, 29)=4.95, p=0.01, Tukey post-hoc test). The mice were more active at night than during the day, consistent with their nocturnal nature (Figure 2A). The mice spent about ∼30% of the night and about ∼10% of the day being active. Thus, the mice were active for a minority of the time not only in their rest phase (day), but also in their active phase (night). Moreover, in the rest phase, the mice were not inactive, but rather active for 10% of the time.

**Figure 2:**
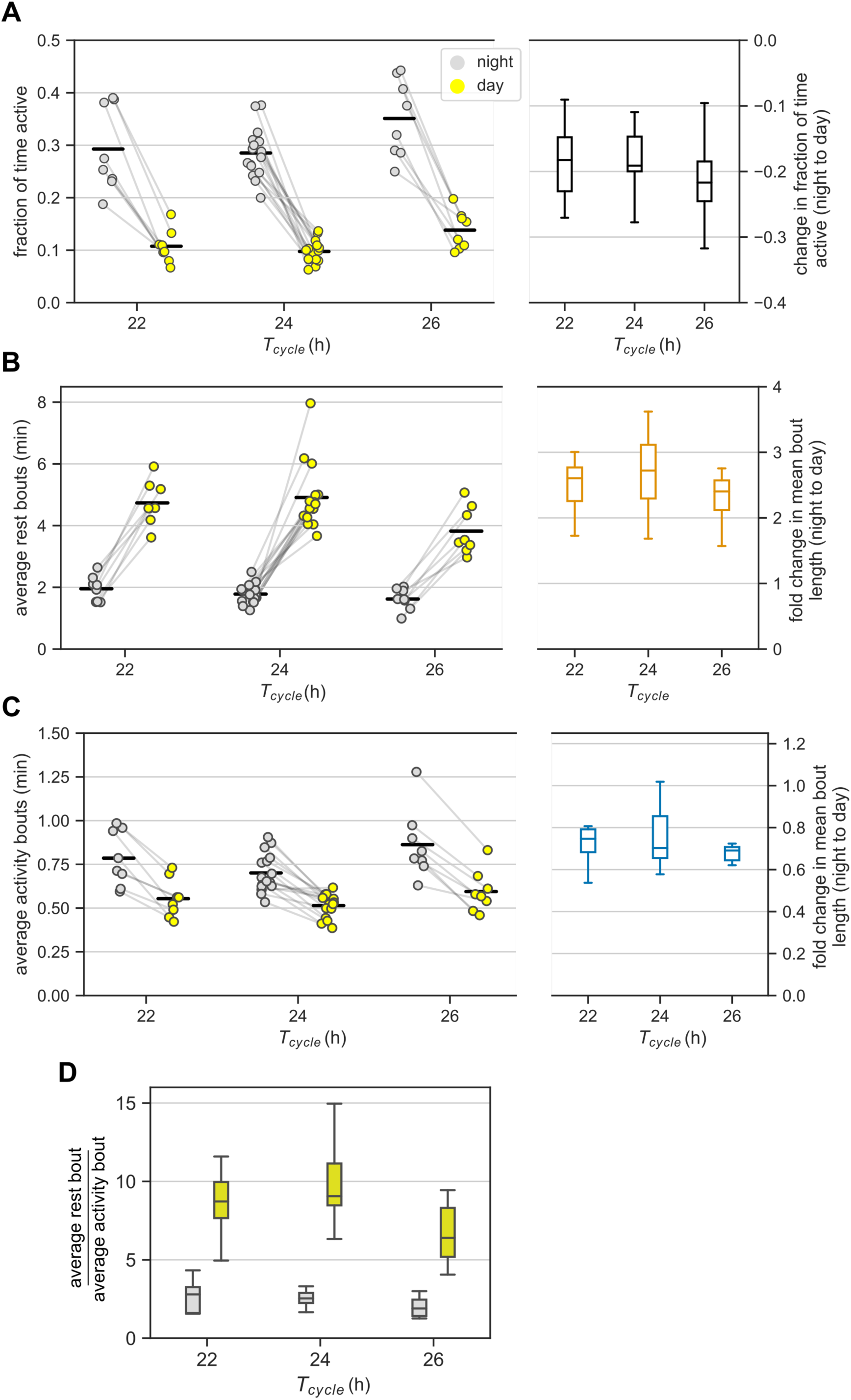
Ultradian behavior under LD cycles: (A) Average activity of mice in the day and night under different LD cycles. Average activity is the fraction of bins with activity during an interval of day or night. Each animal contributed two points (one each for the day and the night) that are connected. The changes in the average activity from night to day for different T-cycles are shown in the boxplot. (B, C) The average bout lengths for rest (B) and activity (C) estimated by the model in the day and the night under different T-cycles. The pair of points (one each for day and night) contributed by each animal is connected. The fold-change in mean bout length from night to day is provided as boxplots on the right. (E) The relative lengths of mean rest and mean activity bouts in the night and day for different T-cycles. The same data in (B) and (C) are visualized differently in (E). Color-coding is maintained throughout the figure. Horizontal black bars in the scatter plots represent mean of the values in that column.

The night to day change in average activity was similar among the three T-cycles. Specifically, the average activity during the day was one-third of the levels during the night for all three T-cycles (Figure 2A). The fold reduction of 0.38, 0.35, and 0.40 for the 22-, 24, and 26-hour T-cycles had 95% confidence intervals (CI) of [0.31, 0.45], [0.31, 0.40] and [0.33, 0.47], respectively. Despite the higher average activity under the 26-hour T-cycle, the night to day change in the 26-hour T-cycle was indistinguishable from the night to day change in the other two T-cycles.

### 3.3 Activity and rest bouts are inversely regulated during the day and night

Next, we examined whether the average duration of the activity and rest bouts were different between the day (i.e., the resting phase) and the night (i.e., the active phase) for the three different T-cycles.

We observed higher activity during the night than during the day (Figure 2A). This could result from three different scenarios: (i) rest bouts are longer during the day than during the night, but activity bouts are unchanged between day and night (ii) activity bouts are longer during the night than during the day, but rest bouts are unchanged between day and night (iii) rest bouts are shortened and activity bouts are lengthened from day to night. In this section, we identify the scenario that is most consistent with the data.

According to our analysis, on average, the rest bouts were longer during the day compared to during the night under all three T-cycles (Figure 2B). Average bouts of rest during the day were 2.75-fold longer during the night (CI: [2.44, 3.09]) with no significant differences across the three T-cycles; fold-change for the 22-hour and the 26-hour T-cycles relative to the 24-hour T-cycle had CIs of [0.94, 1.27] and [0.78, 1.05].

On the other hand, the activity bouts on average were shorter during the day compared to during the night under all three T-cycles (Figure 2C). Average activity bouts were 0.74-fold shorter during the day compared to during the night with only 26-hour T-cycle having a slightly larger decrease in the activity bout length; fold-change for 22-hour and 26-hour T-cycles relative to 24-hour T-cycle had CIs of [0.96, 1.29] and [1.05, 1.41].

Thus, both rest and activity bouts are indeed regulated reciprocally between day and night.

### 3.4 Day-night changes in rest bouts dominate day-night changes in activity bouts

This section compares the relative durations of rest and activity bouts during the day and during the night.

We observed that the average rest bout was always longer than the average activity bout (Figure 2B,C, ratio_rest/activity_=2.53, CI: [2.25, 2.86]). This agrees with mice having more rest than activity (average activity < 0.5) during both day and night (Figure 2A). The average rest bout was about twice as long as the average activity bout in the night (ratio_rest/activity_ = 2.28, CI: [2.08, 2.49], Figure 2D). The ratio increased to about eight-fold in the day (ratio_rest/activity_ = 8.06, CI: [7.37, 8.81]) and was significantly greater under the 24-hour T-cycle (ratio = 1.17, CI: [1.05, 1.31]).

It appears therefore that the average rest bout is always longer than the average activity bout and the absolute day-night change in the rest bouts is also greater than the absolute day-night change in the activity bouts (Figure 2B,C,D). We therefore hypothesize that the day-night changes in rest bouts dominate the day-night changes in activity bouts. Quantifying the day-night change in the number of bouts can help test this hypothesis.

Given that “rest” is defined as the lack of activity, bouts of rest and bouts of activity always alternate (Figure 1B,C); therefore, the number of rest and activity bouts in any given time interval is equal (or differs by no more than one). As a result, we only report the total number of bouts in an interval. If rest bouts were to dominate the day-night change, then the number of bouts would be expected to be higher during the night than during the day (rest bouts are shorter during the night). If, on the other hand, the activity bouts dominate, then the number of bouts would be expected to be lower during the night than during the day. Since the total number of bouts during the night was higher than during the day (Figure S3A), we conclude that rest bouts rather than activity bouts dominate the day-night change in activity.

### 3.5 Mice in DD are less active than in LD due to reduced activity during the subjective night

In LD cycles, both light and the central pacemaker influence behavioral activity, while DD conditions allow us to study behavior without the influence of light. This section compares ultradian behavior in DD with LD in order to distinguish between the effect of the circadian system and light.

In DD, mice had lower average activity overall, particularly during the subjective night. Mice in DD were about 20% less active than mice in LD (Figure S2, *F*(1, 47) = 12.33, p < .001, ratio_DD/LD_ = 0.79); we pooled data from all three T-cycles in Figure 2 into the LD group. In DD, mice were active for about 20% of the time during the subjective night and about 12% of the time during the subjective day. Thus, a difference in activity between night and day also existed in DD (Figure 3A, ratio_light/dark_ = 0.59, CI: [0.53, 0.67])), but was considerably smaller than in LD (ratio_DD/LD_=0.60, CI: [0.52, 0.70]). Interestingly, this difference resulted only from reduced activity during the subjective night in DD (ratio_DD/LD_ during the subjective day = 1.12, CI: [0.96, 1.29]; ratio_DD/LD_ during the subjective night = 0.67, CI: [0.58, 0.78]).

**Figure 3:**
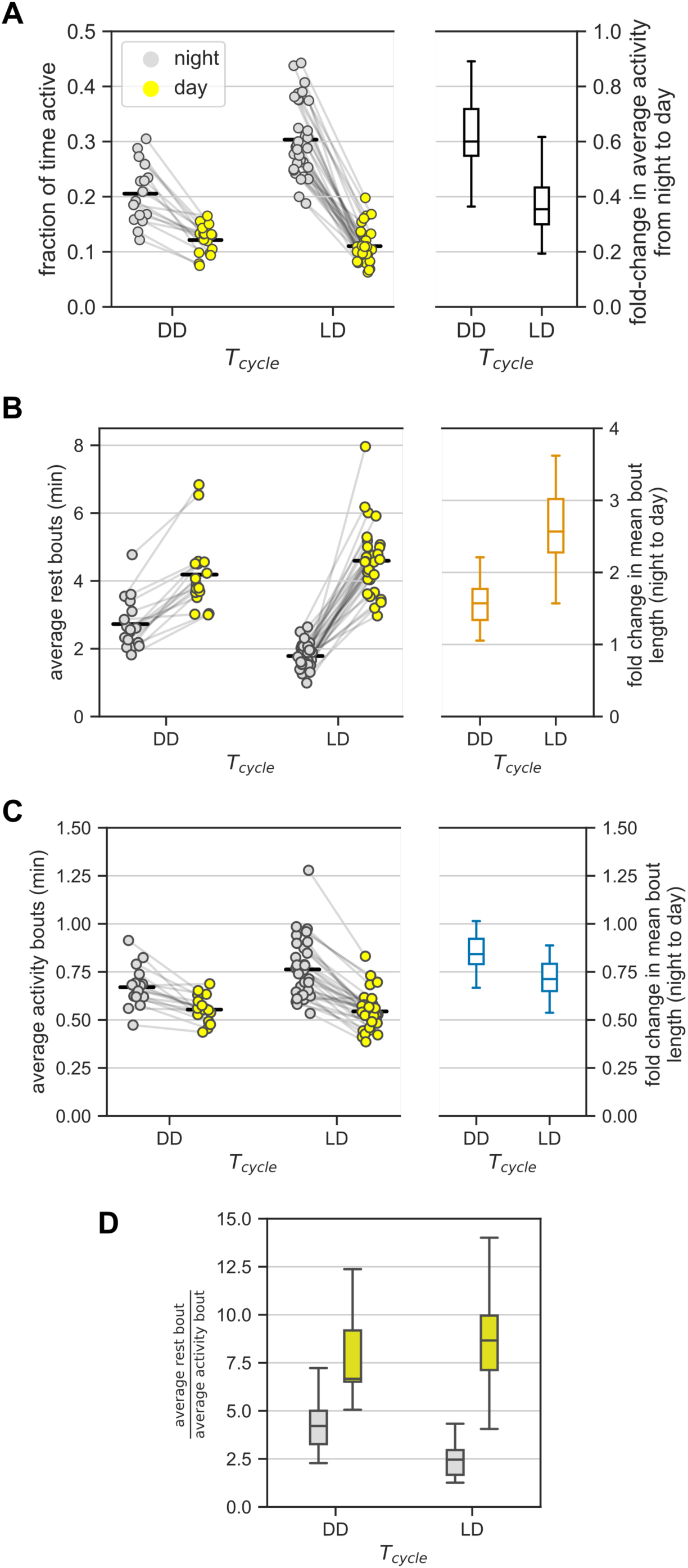
Ultradian behavior under constant darkness (DD). (A) Average activity of mice (measured as the fraction of active bins) during the day and during the night under DD and LD cycles. In DD, day and night refer to *subjective* day and *subjective* night, respectively. The LD group consists of data from the three T-cycles. Each animal contributed two points, one each to the day and the night groups – pairing is denoted by gray lines. Fold-change in the average activity between night and day is presented as a boxplot. (B, C) The model-based estimates of the mean rest (B) and mean activity (C) bout lengths in the (subjective) night and (subjective) day in DD and LD. Data points from the same mice are connected by gray lines. The bout length averaged over all individuals is marked with black bars. Boxplots show the fold-change in bout lengths from night to day. (D) The relative lengths of the mean rest and mean activity bouts in the (subjective) day and (subjective) night in DD and LD (a different representation of data in (B) and (C)).

### 3.6 Day-night changes in bout lengths persisted in DD, but were moderated

This section continues the comparison between DD and LD cycles with a focus on the model-based estimates of mean bout lengths.

Mean rest bout and mean activity bout lengths changed inversely between subjective day and subjective night also in DD (Figure 3B,C). The mean length of the rest bouts increased 1.5-fold from the subjective night to the subjective day (Figure 3B, Table 1), whereas the mean length of the activity bouts decreased by 20% from the subjective night to the subjective day (Figure 3C, Table 1).

**Table 1:** Mean and 95% confidence intervals (CI) for the average bout lengths under DD and LD cycles. The relative values of the bouts lengths are included as ratios (including their respective CIs)

|  | DD |  |  |  | LD |  |  |
| --- | --- | --- | --- | --- | --- | --- | --- |
| | subjective<br>night | subjective<br>day | $\frac{\text{subj. day}}{\text{subj. night}}$ | | night | day | $\frac{\text{day}}{\text{night}}$ |
| activity<br>bout | 0.66 min<br>[0.60, 0.73] | 0.55 min<br>[0.50, 0.60] | 0.83<br>[0.73, 0.94] |  | 0.75 min<br>[0.70, 0.80] | 0.54 min<br>[0.50, 0.57] | 0.71<br>[0.65, 0.79] |
| rest bout | 2.64 min<br>[2.41, 2.89] | 4.08 min<br>[3.72, 4.47] | 1.54<br>[1.36, 1.75] |  | 1.75 min<br>[1.64, 1.87] | 4.50 min<br>[4.21, 4.81] | 2.57<br>[2.35, 2.82] |
| $\frac{\text{rest}}{\text{activity}}$ | 3.99<br>[3.51, 4.52] | 7.42<br>[6.54, 8.41] | 1.86<br>[1.56, 2.23] | | 2.34<br>[2.13, 2.56] | 8.39<br>[7.65, 9.19] | 3.59<br>[3.15, 4.08] |

This day-night change in the mean length of the rest bouts in DD was smaller in comparison to the corresponding change in LD (Figure 3B, ratio_DD/LD_=0.6, CI: [0.51, 0.70]). However, the day-night change in the mean length of the activity bouts was statistically indistinguishable between LD and DD conditions (Figure 3C, ratio_DD/LD_=0.86, CI: [0.74, 1.01]). Thus, day-night changes in the rest bout lengths, but not in the activity bout lengths, were moderated in DD.

The mean rest bouts were longer than the mean activity bouts during both the subjective day and the subjective night in DD (Figure 3D). The day-night changes in the rest bout lengths and the activity bout lengths result in different ratios of mean rest bout and mean activity bout lengths between subjective day and subjective night. The ratio of rest bout lengths to activity bout lengths during the night was significantly greater under DD than under a LD cycle (Figure 3D, Table 1). However, the ratio of rest bout lengths to activity bout lengths during the day was similar in DD and LD (Table 1). As a result, the ratio of rest bout lengths to activity bout lengths varied less in DD than in LD.

Day-night changes in the rest bouts rather than in the activity bouts predominantly contributed to the day-night changes in activity. The number of total bouts was higher during the subjective night compared to the subjective day, which coincides with the shorter rest bouts during the subjective night (Figure 3B). However, the day-night difference in the total number of bouts was smaller in DD than in LD (Figure S3B).

### 3.7 Model underestimates the number of very short and very long rest bouts

The probabilistic model fitting ensures that the mean bout lengths in the behavioral data and the model are identical (fitting probability parameters is tantamount to fitting the means). In this section, the observed distribution of the rest and activity bout lengths is contrasted against the distribution predicted by the model.

The model-derived rest bout length distribution deviates from the observed distribution. Under the probabilistic model (Figure 1A), bout lengths follow a geometric distribution with a mean given by the model parameters. The model predicts fewer extremely short and extremely long rest bouts than those observed in the data (Figure 4). Nevertheless, the activity bout distribution in the data closely matched the predicted distribution. This predicted rest bout distribution consistently differed from the data across the three T-cycles and constant darkness.

**Figure 4:**
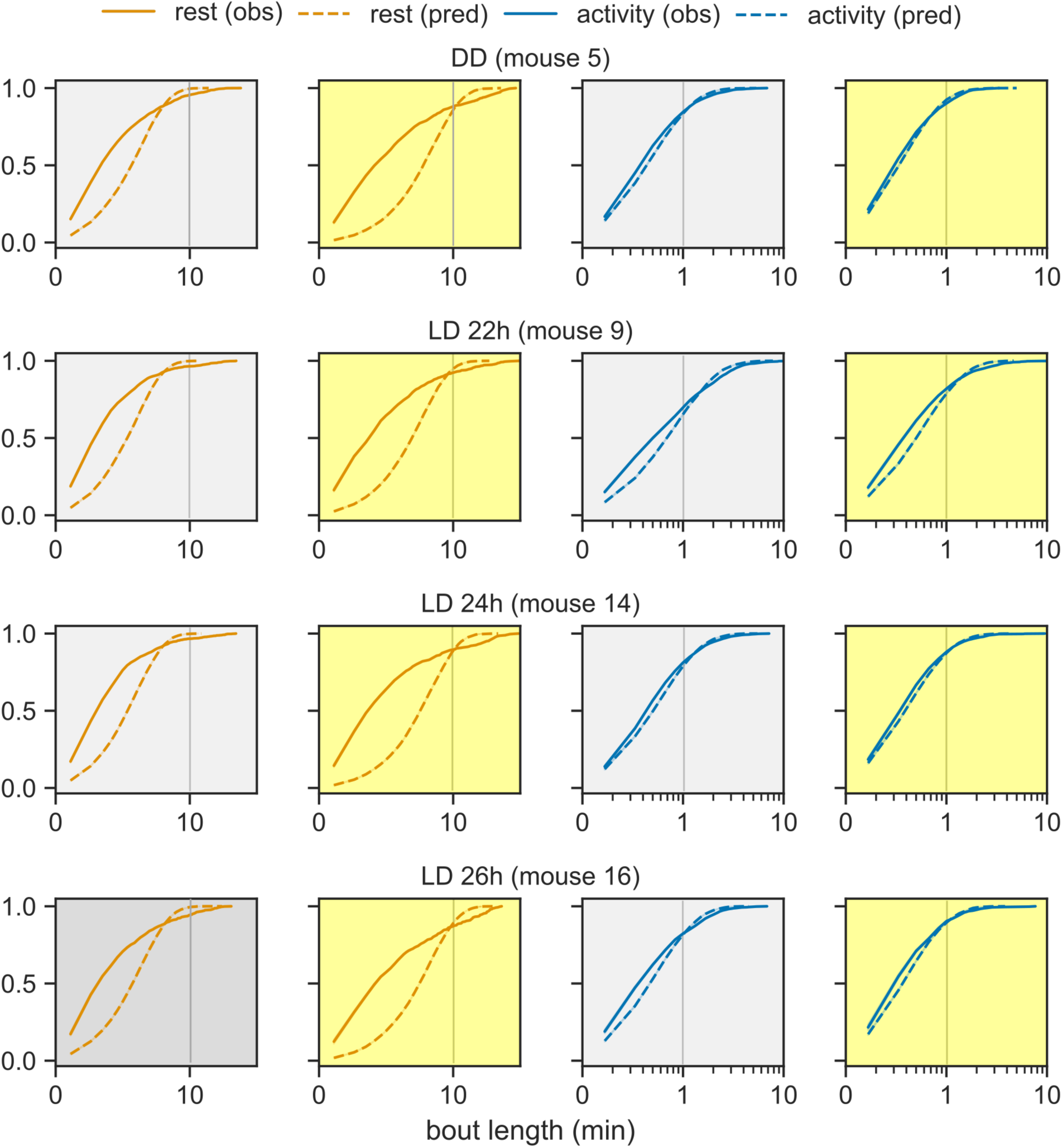
Distribution of bout lengths of activity and rest. The observed distribution of bout lengths (solid lines) is compared against the distribution predicted by the model (dashed lines). The rest and activity bouts in the day and in the night are plotted in separate panels for one representative animal from each group: LD cycles of 22h, 24h and 26h and DD. The x-axis is plotted in logarithmic scale in order to see the very short and very long bouts in one graph. The color-coding from earlier figures is maintained.

## 4 Discussion

The central pacemaker in mammals contributes to the daily rhythmic patterns of behavioral activity. This paper set out to test whether behavioral activity is also regulated within the 24h circadian cycle. Using data on spontaneous behavioral activity of mice under LD and DD, we quantified activity in terms of rest and activity bouts in the day and in the night using a probabilistic model. We observed day-night differences in the average bouts lengths of rest and activity under LD that persisted also under DD. The probabilistic model was able to exploit the structure in the behavioral activity to accurately capture the distribution of bouts (with the exception of extremely short and extremely long rest bouts). The model is evidence that (a) behavioral activity is indeed regulated at the level of bouts and (b) these bouts are under the control of both light input and the circadian system.

The main analytical contribution of this work is the probabilistic model of mouse behavior. A probabilistic, as opposed to a deterministic, model is necessary, since there is clearly large intra-individual (across the length of the recordings) and inter-individual variability even in isogenic mice kept under identical conditions. A simpler model would have a single probability parameter defining the switch between rest and activity and vice-versa (i.e., *α* = *β*). But, the stark difference in activity between the day and night makes this simpler model inadequate. The model we propose is thus the simplest (non-trivial) model of mouse behavior in the ultradian timescale. Such Markov models are well established in many fields including genomics (Pardoux, 2010) and sleep research (Kemp and Kamphuisen, 1986; Stephenson et al., 2013).

Conveniently, the model directly relates to behavior of mice at the level of bouts. Early studies qualitatively described the organization of behavior in small mammals into characteristic bouts (Kavanau and Rischer, 1968; Davis and Menaker, 1980). Penev et al. (1997) studied temporal patterns of behavior in hamsters and showed increased fragmentation with age. Farajnia et al. (2012) showed similar fragmentation in aged mice. Ultradian periodicity of behavior in mammals also manifests as rhythmic consolidated bouts of activity. Ultradian rhythms are observed under natural conditions in the common vole (Gerkema and van der Leest, 1991) and mice (Dowse et al., 2010), and after surgical or genetic manipulation of the clock in rats and mice (Vitaterna et al., 1994; Horst et al., 1999; Blum et al., 2014).

Mice were inactive for a majority of the day and the night, but with significant activity even during the day. Nevertheless, the mice showed more activity in the night versus the day. Spontaneous behavior (measured using passive IR sensors) is not as clearly segregated into an active phase and a rest phase as is wheel-running activity (Schwartz and Zimmerman, 1990). The mean rest bouts were shortened and the mean activity bouts lengthened in the night relative to the day in the 22-hour, 24-hour and 26-hour T-cycles. The different T-cycles affect clock function under the entrained conditions studied here. Since the day-night differences in bout lengths were unaltered across T-cycles, we conclude that light regulates the length of rest and activity bouts independent of the central clock. To fully support the latter conclusion, we may study ultradian behavioral regulation in animals under short and long photoperiod, as an additional modifier of clock function.

We observed that, on average, rest bouts were always longer than activity bouts. Moreover, the day-night changes in mean rest bout lengths were about two-fold larger than the changes in mean activity bout lengths in LD. Taken together, the day-night changes in rest bouts (in minutes) is significantly larger than day-night changes in activity bouts. Therefore, we conclude that regulation of rest bouts predominantly underlies the differences between the active and rest phases. If this were true, we would expect higher number of bouts (rest and activity) in the night compared to the day. This is indeed the case. Thus, the LD environment regulates rest bouts rather than activity bouts over the 24h cycle.

Under a LD cycle, both light and the central clock affect behavioral activity. To determine the effects of the central clock on behavior, animals are routinely exposed to constant darkness, and in the absence of all potential time cues. In DD, mice continued to show the same qualitative changes in mean bout lengths between subjective day and subjective night. The persistence of bout regulation in DD between subjective day and night suggests that the central clock also regulates bout length.

The day-night difference in activity and rest bout duration was larger in LD cycles than in DD. Given the enhanced difference between the day and night under LD cycles, we conclude that environmental light cycles reinforce the SCN effect on bout regulation. In other words, exposure to LD cycles increases the “amplitude” of circadian regulation of bouts. Interestingly, the amplitude of the circadian rhythm is also increased under LD as compared to DD conditions. This is the case both at the level of behavioral activity and also at the level of SCN electrical discharge rate (Coomans et al., 2013). It is possible therefore, that even the influence of light on ultradian behavior involves the SCN. At least, and given our results obtained from DD conditions, we suggest that the SCN is a node in the central regulation of ultradian behavioral activity, and is a regulator of the duration of ultradian bouts. Thus, in the absence of the SCN, ultradian bouts will still be present, but the day-night difference in their duration will be completely lost. This interpretation is in line with the ongoing presence of ultradian behavioral rhythmicity in transgenic clockless animals (Vitaterna et al., 1994; Horst et al., 1999; Blum et al., 2014), as well as in voles with SCN lesions (Gerkema et al., 1990).

Surprising, the Markov model predicted well the behavior at the next 10s bin based on the current bin and a coin toss. We also confirmed explicitly the validity of the Markov assumption for our behavioral data. The rest and activity states in the Markov model have positive (auto-enforcing) feedback loops (Figure 1). When the strength of this positive feedback is large (>0.5, which is the case for all fits in this study), then the Markov model shows inertia, i.e., a tendency to remain at the state it is in. Occasionally, behavior breaks out of this state and switches to the alternative state. We found that this principle applies well to the ultradian regulation of rest and activity. The parameters describing the duration in a state are apparently under the control of environmental light and the central clock. The model is analogous to the proposed “flip-flop” switch between sleep and wakefulness, where various neuronal inputs regulate the balance between the two states (Saper et al., 2001). It is very likely that other brain areas are also involved in the underlying circuitry, and for instance, there is good evidence for the role of dopamine (Blum et al., 2014).

The Markov property is manifested as a geometric distribution of bout lengths, where short bouts are more common than long bouts. While the model predicted the distribution of activity bouts accurately, it underestimated both the number of very short and very long rest bouts. Nevertheless, our model is elegant in its simplicity in that it captures most features of murine behavior with some exceptions. In fact, there are multiple reports showing that especially rest bouts do not follow an exponential distribution. The first quantitative analysis of bouts (Penev et al., 1997) already proposed that rest bouts were of two types: short bouts within an activity bout or long bouts between activity bouts. More generally, rest bout lengths appear to follow a power-law (heavy-tailed) distribution in mice, humans and fruit flies (Nakamura et al., 2008; Cascallares et al., 2018) that breakdown under certain pathologies. The rest bout distribution in this study did not show power-law characteristics (not shown). The deviation of the rest bout distribution might be due to the presence of different types of rest bouts, such as a “pause” in activity explaining the very short bouts, and the existence of other processes regulating rest bouts, such as homeostatic sleep drive.

Our conclusions must be viewed in the context of our analysis methodology. The model treats behavior within the dark and light phases as homogeneous, which is not often the case (Houben et al., 2014). The simplified data capture the duration of activity, but ignore the intensity of activity. Thus, there could be differences in intensity of activity across LD cycles or between DD and LD, which our analysis overlooked. All these limitations provide interesting avenues for further study. Finally, the behavioral activity was collected in 10s bins and although bout lengths were of the order of minutes, the effect of bin size on the results cannot be excluded.

## 7 Author Contributions

BA conceived and designed the study, performed the analysis and wrote the manuscript; JHM acquired the data and critically revised the manuscript; BA and JHM interpreted the data. All authors contributed to manuscript revision, and read and approved the submitted version.

## 9 Acknowledgments

The authors thank Hanspeter Herzel and Achim Kramer for useful discussions and Jos H. T. Rohling for compiling data from prior studies.

