## Supplemental Figures for "Ultradian regulation of rest and activity bouts in mice"

### *Supplementary Material*

#### 1 Supplementary Figures

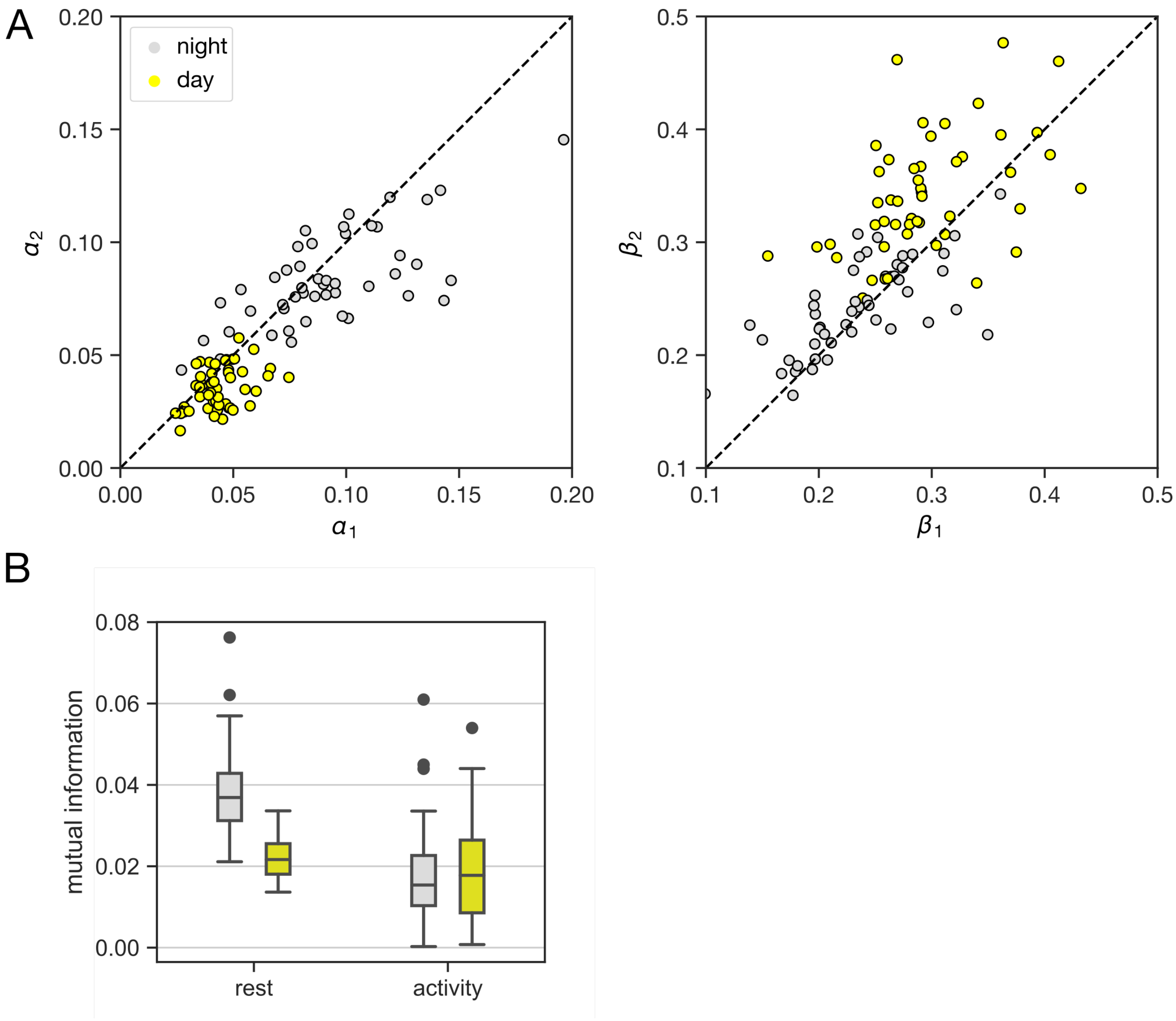


**Supplementary Figure 1.** Robustness and consistency of the model fits. (A) The comparison of the model probability parameters $\alpha$ and $\beta$ estimated from the first half of the data record (x-axis) against the second half (y-axis). Two points in each figure belong to one mouse – the pairing is not shown for clarity. (B) The empirical mutual information (MI) between behavioral state in the next 10s bin and the last 10s conditioned on the current behavioral state (on the x-axis). The MI was computed for the day and night separately. Data from LD cycles and DD were pooled for (A) and (B).

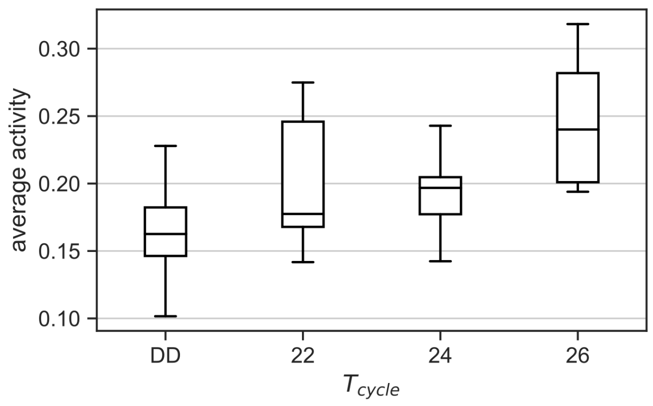


**Supplementary Figure 2:** The average activity under DD and different T-cycles. The average activity of a mouse is the fraction of bins over the entire length of the recording in which the mouse was active. The box-plot represents the estimates of average activity for multiple mice under each condition.

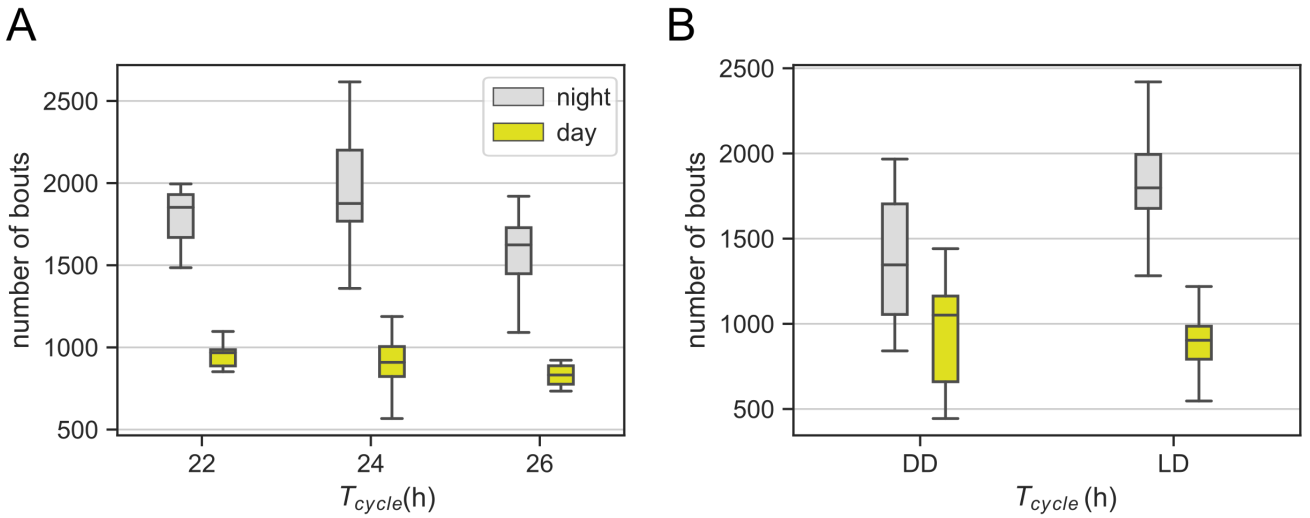


**Supplementary Figure 3.** The total number of bouts during the day and during the night. (A) The variation in the total number of bouts in the day and night for different T-cycles. (B) The total number of bouts in the (subjective) day and (subjective) night in DD versus LD conditions. Since rest is defined as the lack of activity, the number of rest and active bouts are equal (or can differ by at most one). Therefore, the number of both rest and active bouts is a half of the numbers in (A) and (B).
